# Barnaba: Software for Analysis of Nucleic Acids Structures and Trajectories

**DOI:** 10.1101/345678

**Authors:** Sandro Bottaro, Giovanni Bussi, Giovanni Pinamonti, Sabine Reißer, Wouter Boomsma, Kresten Lindorff-Larsen

## Abstract

RNA molecules are highly dynamic systems characterized by a complex interplay between sequence, structure, dynamics, and function. Molecular simulations can potentially provide powerful insights into the nature of these relationships. The analysis of structures and molecular trajectories of nucleic acids can be non-trivial because it requires processing very high-dimensional data that are not easy to visualize and interpret.

Here we introduce Barnaba, a Python library aimed at facilitating the analysis of nucleic acids structures and molecular simulations. The software consists of a variety of analysis tools that allow the user to i) calculate distances between three-dimensional structures using different metrics, ii) back-calculate experimental data from three-dimensional structures, iii) perform cluster analysis and dimensionality reductions, iv) search three-dimensional motifs in PDB structures and trajectories and v) construct elastic network models (ENM) for nucleic acids and nucleic acids-protein complexes.

In addition, Barnaba makes it possible to calculate torsion angles, pucker conformations and to detect base-pairing/base-stacking interactions. Barnaba produces graphics that conveniently visualize both extended secondary structure and dynamics for a set of molecular conformations. The software is available as a command-line tool as well as a library, and supports a variety of file formats such as PDB, dcd and xtc files. Source code, documentation and examples are freely available at https://github.com/srnas/barnaba under GNU GPLv3 license.

## Introduction

Despite their simple four-letters alphabet, RNA molecules can adopt amazingly complex three-dimensional architectures. RNA structure is often described in terms of few, simple degrees of freedom such as backbone torsion angles, sugar puckering, base-base interactions, and helical parameters ***Dickerson*** (***1989***); ***Leontis and Westhof*** (***2001***); ***Richardson et al.*** (***2008***). Given a known three-dimensional structure, the calculation of these properties can be accurately performed using available tools such as MC-annotate ***Gendron et al.*** (***2001***), 3DNA ***Lu and Olson*** (***2008***), fr3D ***Sarver et al.*** (***2008***) or DSSR ***Lu et al.*** (***2015***). These software packages allow for a detailed description of experimentally-derived RNA structures, but are less suitable for analyzing and comparing large numbers of three-dimensional conformations.

The importance of large-scale analysis tools is critical when considering that many RNA molecules are not static, but highly dynamic entities, and multiple conformations are required to describe their properties. In molecular dynamics (MD) simulations ***Šponer et al.*** (***2018***), for example, it is often necessary to analyze several hundreds of thousands of structures. The analysis and comparison of results from structure-prediction algorithms poses similar challenges ***Dawson and Bujnicki*** (***2016***); ***Magnus*** (***2016***); ***Miao et al.*** (***2017***). In order to rationalize and generate scientific insights, it is therefore fundamental to employ specific analysis and visualization tools that can handle such highly-dimensional data. This need has been long recognized in the field of protein simulations, leading to the development of several software packages for the analysis of MD trajectories ***Michaud-Agrawal et al.*** (***2011***); ***McGibbon et al.*** (***2015***); ***Tiberti et al.*** (***2015***). While these software can be in principle used to analyze generic simulations, they do not support the calculation of nucleic-acids-specific quantities out of the box. Notable exceptions are CPPTRAJ ***Roe and Cheatham III*** (***2013***), and the driver tool in PLUMED ***Tribello et al.*** (***2014***), that support the calculation of nucleic acids structural properties, among other features.

A limited number of software packages have been developed with the main purpose of analyzing simulations of nucleic acids. Curves+ ***Lavery et al.*** (***2009***) calculates parameters in DNA/RNA double helices as well as torsion backbone angles. *do_x_*_3*dna*_ ***Kumar and Grubmüller*** (***2015***) extends the capability of the 3DNA package to calculate several base-pairs/helical parameters and torsion angles from GROMACS ***Abraham et al.*** (***2015***) trajectories. The detection of hydrogen bonds/stacking in simulations and the identification of motifs such as helices, junctions, loops, and pseudoknots can be performed using the Motif Identifier for Nucleic acids Trajectory (MINT) software ***Górska et al.*** (***2015***).

Here we present Barnaba, a Python library to analyze nucleic acids structures and trajectories. The library contains routines to calculate various structural parameters (e.g. distances, torsion angles, base-pair and base-stacking detection), to perform dimensionality reduction and clustering, to back-calculate experimental quantities form structures and to construct elastic network models. Barnaba utilizes the capabilities of MDTraj ***McGibbon et al.*** (***2015***) for reading/writing trajectory files, and thus supports many different formats, including PDB, dcd, xtc, and trr.

In this paper we show the capabilities of Barnaba by analyzing a long MD simulation of an RNA stem-loop structure. We first calculate distances from a reference frame. Second, we consider a subset of dihedral angles and compare ^3^*J* scalar couplings calculated from simulations with nuclear magnetic resonance (NMR) data. We then perform a cluster analysis of the trajectory, identifying a number of clusters that are visualized using a dynamic secondary structure representation. Finally, we search for structural motifs similar to cluster centroids in the entire protein data bank (PDB) database. In addition, we show how to construct an elastic network model (ENM) of RNA molecules and protein-nucleic acid complexes with Barnaba, and how to use it to estimate RNA local fluctuations. Source code and documentation are freely available at https://github.com/srnas/barnabaunder GNU GPLv3 license.

## Results

First, we provide a list of tools for the analysis of nucleic acids three-dimensional structures supported in Barnaba. All the calculations can be executed from the command-line, as described in Supplementary Material (SM1). For each functionality, practical examples are provided in Supplementary Material and in the documentation:

1. Calculate the eRMSD ***Bottaro et al.*** (***2014***) between structures. (SM2)
2. Calculate the heavy-atom/backbone-only root mean squared distance (RMSD) after optimal superposition ***Kabsch*** (***1976***) between structures (SM2).
3. Calculate the relative position and orientations between nucleobases (SM3).
4. Identify base-pairing and base-stacking interactions in structures and trajectories (SM4).
5. Calculate backbone, sugar and pseudorotation torsion angles (SM5).
6. Back-calculate ^3^J scalar couplings from structures (SM6).
7. Search for single-stranded and double-stranded three-dimensional motifs within PDB structures or trajectories (SM7-SM8).
8. Extract fragments with a given sequence from PDB structures. This can be useful to investigate the conformational variability of RNA at a fixed sequence or to perform a stop-motion modeling (SMM) analysis ***Bottaro et al.*** (***2016b***) (SM9).
9. Perform cluster analysis of RNA structures using the eRMSD (SM10).
10. Generate “dynamic secondary structure” figures, that display the extended secondary structure, together with the population of each interaction within a collection of three-dimensional structures (SM11).
11. Construct elastic network models (ENM) of RNA molecules and protein-nucleic acid complexes (SM12).
12. Calculate the scoring function eSCORE ***Bottaro et al.*** (***2014***); ***Poblete et al.*** (***2018***) (SM13).

In the following, we present the different features of Barnaba by analyzing a 180*μs* long simulation of an RNA 14-mers with sequence GGCACUUCGGUGCC performed by Tan et al. ***Tan et al.*** (***2018***) using a simulated tempering protocol where the temperature is used as a dynamic variable to enhance sampling. Experimentally, this sequence is known to form an A-form stem composed by 5 consecutive Watson-Crick base pairs, capped by a UUCG tetraloop (Fig. 1A). In order to make the results described in this paper fully reproducible, we provide in supplementary material 14 the jupyter notebooks to conduct the analyses and to produce the 1gures described below.

### RMSD, eRMSD calculation and detection of base-base interactions

We start the analysis by calculating the distance of each frame in the simulation from the reference experimental structure (PDB code 2KOC ***Nozinovic et al.*** (***2010***)) and detecting base-base interactions. Fig.1B shows the time series of heavy-atom root mean squared distance (RMSD) after optimal superposition ***Kabsch*** (***1976***). During this simulation, multiple folding events occur: In line with previous analyses ***Tan et al.*** (***2018***) we thus observe both structures close to the reference as well as unfolded/misfolded ones. We identify the base-base interactions in each frame using the annotation functionality in Barnaba (see Methods). Structures where the stem is completely formed together with the native trans sugar-Watson (tSW) interaction between U6-G9 in the loop are shown in red. Blue points indicate structures in which all base pairs in the stem, but not in the loop, are present. All the other structures are colored in gray. From the histogram in Fig. 1B it can be seen that RMSD < 0.23nm roughly corresponds to native-like structures. A second sharp peak around 0.3nm corresponds to structures in which only the stem is correctly formed. All other conformations have RMSD larger than 0.6nm.

One of the features of Barnaba is the possibility to calculate the eRMSD ***Bottaro et al.*** (***2014***). The eRMSD only considers the relative arrangements between nucleobases in a molecule, and quantifies the differences in the interaction network between two structures. In this perspective, eRMSD is similar to the Interaction Fidelity Network ***Parisien et al.*** (***2009***), that quantifies the discrepancy in the set of base-pairs and base-stacking interactions. The eRMSD, however, is a continuous, symmetric, positive definite metric distance that satisfies the triangular inequality. Additionally, it does not require detection of the interactions (annotation) and is hence particularly well suited for analyzing MD trajectories and unstructured RNA molecules. Fig.1C shows the eRMSD from native for the UUCG simulation. We notice that, similarly to the RMSD case, the histogram displays three main peaks. In this case the correspondence between peaks and structures can be readily identi1ed: when eRMSD*<*0.7 native stem and loop are formed, if 0.7*<*eRMSD*<*1.3, stem is formed but the loop is in a non-native con1guration. Other structures typically have eRMSD>1.3. We observe that the separation between the two main peaks (native structure, red, and native stem, blue) is sharper in Fig.1C, con1rming that eRMSD is more suitable than RMSD to distinguish structures with different base pairings ***Bottaro et al.*** (***2014***).

**Figure 1.**
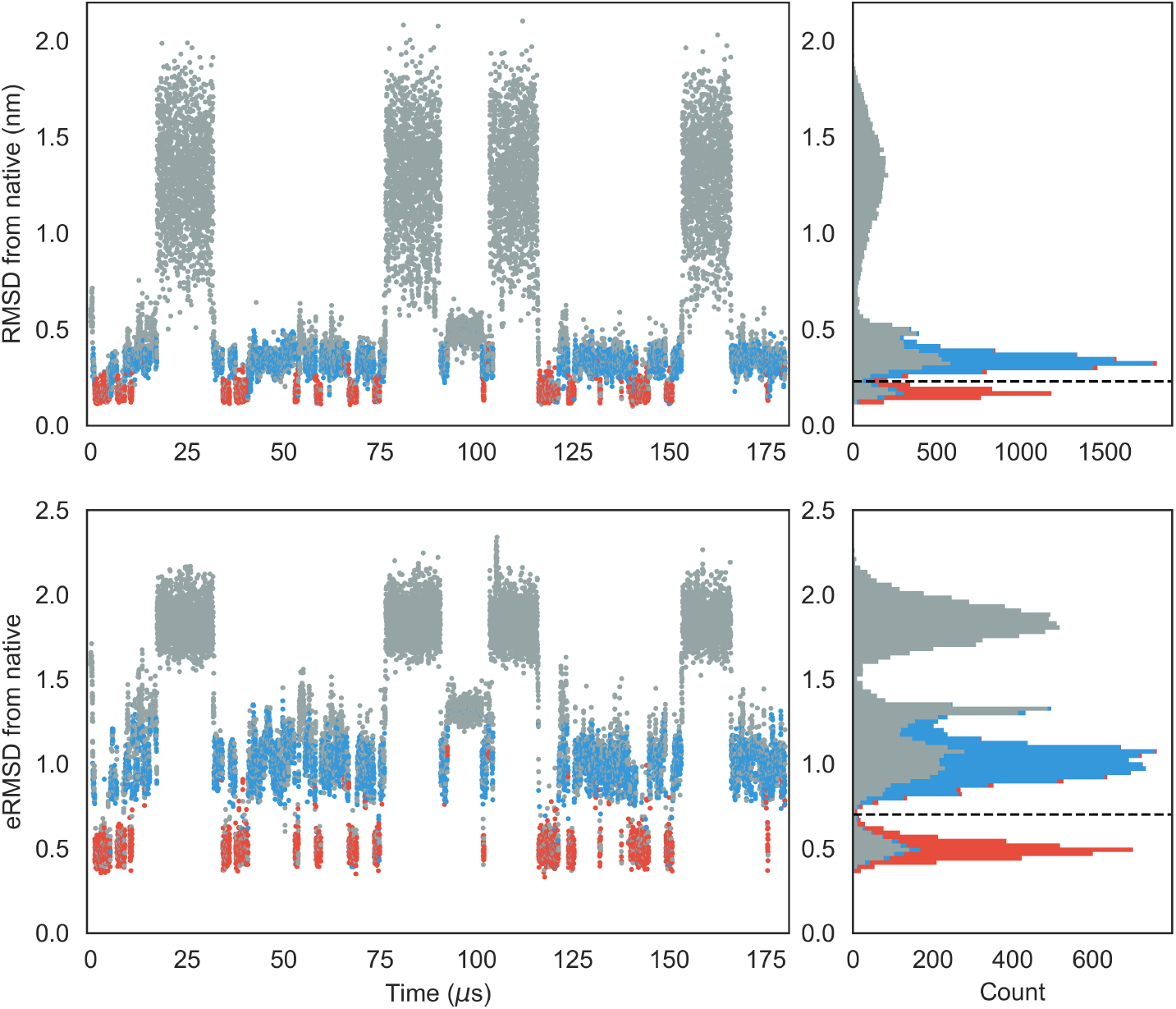
**A**) Extended secondary structure representation of the UUCG stem-loop. Watson-Crick base pairs are shown in blue, trans Sugar-Watson base pair between U6 and G9 is shown in red. **B**) RMSD from native over time of the UUCG simulation. The corresponding histogram is shown in the right panel. The dashed line at RMSD=0.23nm separates native-like from non-native-like structures. The colors indicate the presence of native base-base interactions, as shown in the secondary structure representation. Structures where all Watson-Crick interactions in the stem and the trans Sugar-Watson base pair in loop is formed are shown in red. Blue indicates structures where only the stem is formed. All other conformations are shown in gray. **C**) eRMSD from native structure over time. Color scheme is identical to panel **B**. Dashed line at eRMSD=0.7 separates native-like from non-native conformations.

Note that a signi1cant number of low-RMSD/eRMSD structures lack one or more native base-pair interactions, and are therefore shown in gray. This is because the detection of base-base interactions critically depends on a set of geometrical parameters (e.g. distance, base-base orientation, etc.) that were calibrated on high-resolution structures. The criteria used in Barnaba (as well as the ones employed in other annotation tools) may not always be accurate when considering intermediate states and partially formed interactions that are often observed in molecular simulations ***Lemieux and Major*** (***2002***).

### Transition paths

We now analyze the folding/unfolding paths, in order to understand what is the nature and order of events leading to folding. In particular, we consider the formation of the native base-pairs in the stem and the rotameric state of the *x* angle in G9, that is related to the formation of the Sugar-Watson base pair between G9 and U6. Following Ref. ***Lindorff-Larsen et al.*** (***2011***), we extract the transition paths (TP) from the simulation, resulting in 4 folding and 4 unfolding events. The time evolution for one of the folding events is shown in Fig. 2A-B. In the unfolded state, no base-pairs are formed and *χ* freely fluctuates from *anti* to *syn*. The three base-pairs at the termini form early during the transition path, followed by the other two Watson-Crick base-pairs. When the native state is reached, all native base-pairs are formed, and *χ* is in *syn* conformation.

The order of the events can be quantified by calculating the average presence of base-pair (assuming values of 1=formed or 0=not formed) and the normalized distance from *syn* conformation *q* = 0.5(1 + cos (*χ* − 63°)). Quantities that reach a native-like value (i.e. 1) early during folding have a high value and those that form late get a low value ***Lindorff-Larsen et al.*** (***2011***). In Fig.2C we can see that the Watson-Crick 1-14, 2-13, 3-12 form very early in folding, followed by 4-11 and 3-10. The transition of the *χ* angle to *syn* occurs at a later stage, and folding is finally achieved with the formation of the tSW base-pair.

The TP analysis is here performed for illustrative purposes. In real applications, it is important to take into considerations a number of aspects, such as the quality of the force-field, the assumption that the simulated tempering trajectory is compatible with the real folding pathway, and the employed criteria defining folded/unfolded states ***Lindorff-Larsen et al.*** (***2011***). Note also that this type of analysis is carried out to describe the properties on the energy barrier, while we here describe the properties of the intermediate state.

**Figure 2.**
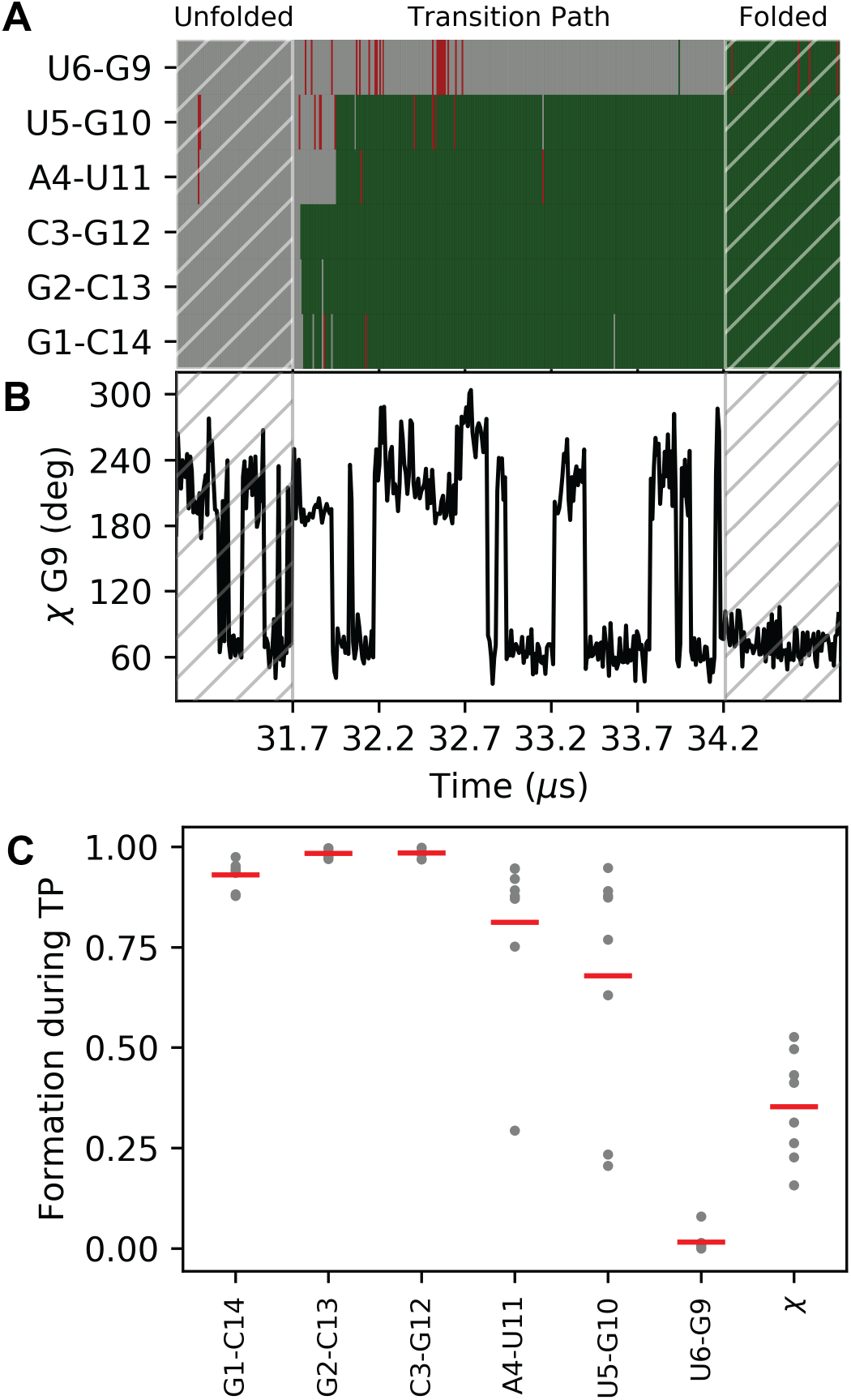
Formation of base-pairs and G9-*χ* angle during RNA folding. A) Formation of the 5 WC-base-pairs over time and of the tSW interaction during one of the folding events. Green indicates that the interaction is formed, gray not formed. Non-native interactions are shown in red. B) Time evolution of the *χ* angle in G9 for the same folding event shown in panel A. C) Order of events relative to the formation of the native base-pairs and transition to *syn*. High values correspond to early formation of the corresponding quantity during a folding event. The average over each TP is shown as a gray dot, and the average over the 8 TPs is shown as a red bar.

### Torsion angle and 3*J* scalar coupling calculations

Another important class of structural parameters is torsion angles. Similarly to other software, Barnaba contains routines to calculate backbone torsion angles (*α*,*β*,*γ*,*∊*,*ζ*,) the glycosidic angle *χ*, and the pseudorotation sugar parameters ***Altona and Sundaralingam*** (***1972***); ***Rao et al.*** (***1981***).

In Fig. 3, left panels, we plot the probability distributions of four angles (*α*,*β*,*γ,δ* and *ϵ*) for three different residues: U6, U7, and G9. We can see from the distribution of *γ* angles that U6 and U7 mainly populate the *gauche*^+^ rotameric state (0° < *γ* ≤ 120°), while G9 significantly populates the *trans* state as well (120° < *γ* ≤ 240°). Different rotameric states can be also seen from the distribution of *8* angles (C2’/C3’-endo) and *ϵ*, that is related to BI/BII states. Here, we consider the same trajectory of the UUCG tetraloops described above and removed all the unfolded structures, i.e. structures with eRMSD from native larger than 1.5 (≈ 6000 out of 20000), because we below compare to experiments under conditions where these are absent.

**Figure 3.**
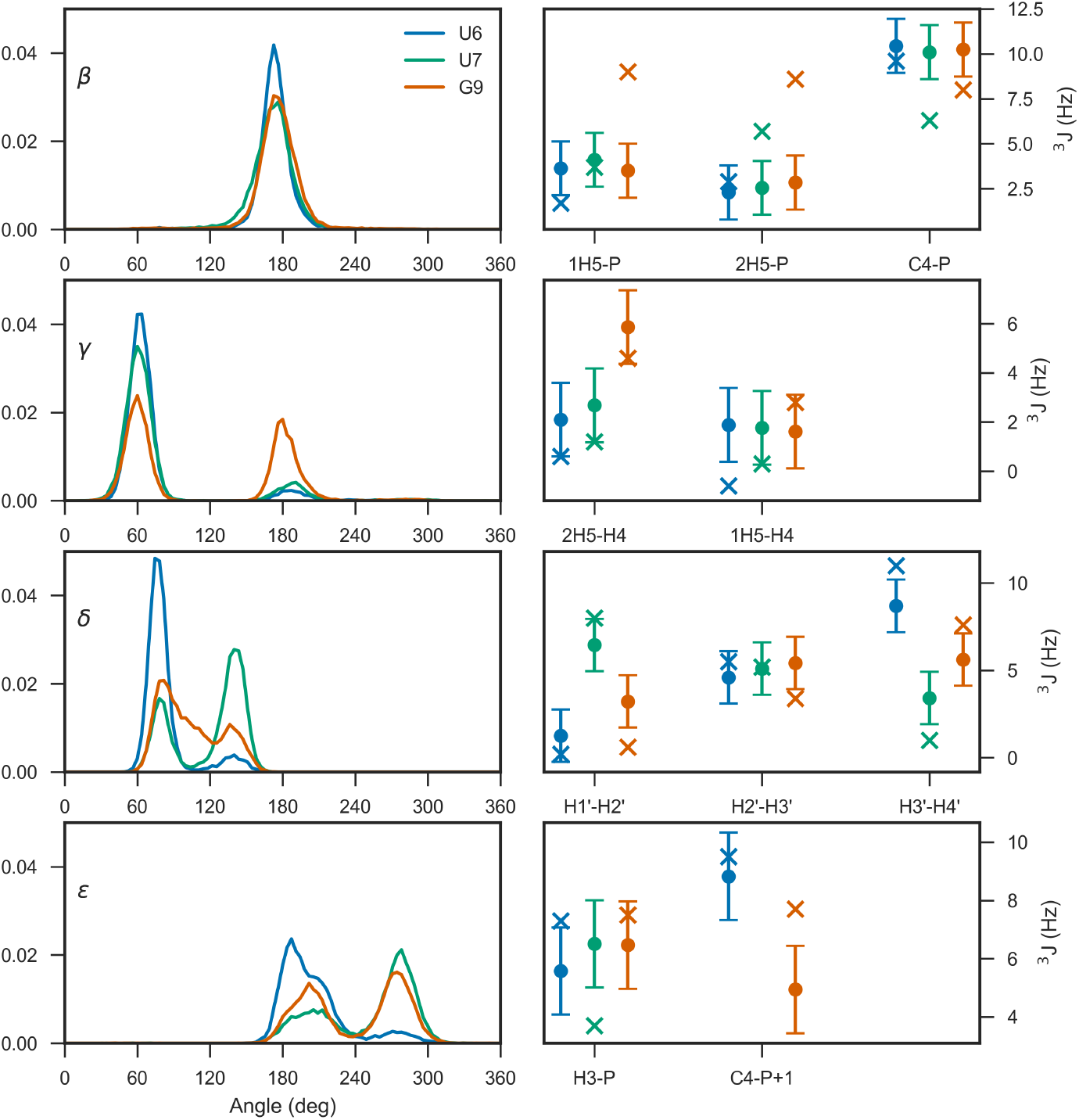
Left panels: Torsion angle distribution for *β*,*γ*,*δ* and *ϵ* in residues U6, U7, and G9. Right panels show the experimental ^3^*J* couplings (crosses) and the calculated value from simulation (dots). The error bars indicate the standard error of the mean calculated over 4 blocks.

In this example we chose these specific torsion angles because their distribution is related to available ^3^J couplings experimental data from nuclear magnetic resonance (NMR) spectroscopy. The magnitude of ^3^J coupling depends on the distance between atoms connected by three bonds, and thus on the corresponding dihedral angle distribution. The dependence between angle *θ* and coupling ^3^*J* can be calculated via Karplus equations ^3^*J* = *A* cos^2^(*θ* + *ϕ*) + *B* cos(*θ* + *ϕ*) + *C*, where *A, B, C* are empirical parameters. Couplings corresponding to different angles can be calculated with Barnaba. H1’-H2’, H2’-H3’, H3’-H4’ (sugar conformation), H5’-P, H5”-P, C4-P (*β*), H4’-H5’, H4’-H5” (*γ*), H3-P(+1), C4-P(+1) (*ϵ*), H1’-C8/C6, and H1’-C4/C2 (*χ*). The complete list of Karplus parameters is reported in the Methods section, and may be changed within Barnaba.

Fig. 3, right panels, show the back-calculated average ^3^*J* couplings and the corresponding experimental value reported in ***Nozinovic et al.*** (***2010***). Note that in some cases experiments and simulations do not agree: this is because the simulation was performed at different temperatures using a simulated tempering protocol, and therefore the comparison between simulations and experiments is here made for illustrative purposes only. Significant discrepancies could originate from errors introduced by the Karplus equations, that can be as large as 2Hz ***Bottaro et al.*** (***2018***).

### Cluster analysis

The structures within a trajectory can be grouped into clusters of mutually similar conformations, to understand which different states are visited and how often. For clustering we use the DBSCAN ***Ester et al.*** (***1996***) algorithm with *ε* = 0.12 and min samples=70 ***Bottaro and Lindorff-Larsen*** (***2017***). As in the previous example, structures with eRMSD > 1.5 from native are discarded. Figure 4A shows the trajectory projected onto the first two components of a principal component analysis done on the collection of **G**-vectors ***Bottaro and Lindorff-Larsen*** (***2017***). Circles show the resulting 9 clusters, whose radius is proportional to the square root of their size. The 5500 structures (40%) that were not assigned to any cluster are shown as gray dots. For each cluster we identify its centroid, here defined as the structure with the lowest average distance from all other cluster members.

Ideally, clusters should be compact enough so that the centroid can be considered as a representative structure. This information is shown in the box-plot in Fig. 4B, that reports the distances (eRMSD and RMSD, as labeled) between centroids and cluster members. At the same time, structures within clusters are not all identical to one another. In order to visualize the intra-cluster variability we have found it useful to introduce a “dynamic secondary structure” representation. In essence, we detect base-stacking/base-pair interactions in all structures within a cluster, and calculate the fraction of frames in which each interaction is present. The population of each interaction is shown by coloring the extended secondary structure representation (Fig.4C). This representation has some analogy with the “dot plot” representation used to display secondary structure ensembles obtained using nearest neighbor models, that reports the predicted probability of individual base pairs ***Jacobson and Zuker*** (***1993***). We can see that the first three clusters correspond to three different tetraloop structures. In cluster 1, the U6-G9 tSW base pair is present, together with the U6-C8 stacking typical of the native UUCG tetraloop structure. In cluster 2, no U6-G9 base pair is present, while in cluster 3 we observe stacking between U6-U7-C8-G9, as also described in the next section. In all clusters the population of the terminal base pairs and stacking is lower than one, indicating the presence of base fraying.

In our experience, cluster analysis is useful to understand and visualize qualitatively the different type of structures in a simulation. In many practical cases, however, the number of clusters and their population may differ depending on the employed clustering algorithm and associated parameters. Clustering may not even be meaningful when considering highly unstructured systems such as long single-stranded nucleic acids lacking secondary structures ***Chen et al.*** (***2012***).

### Motif search

Barnaba can be used to search for structural motifs in a PDB file or trajectory using the eRMSD distance. In the following example, we illustrate this feature by taking the centroids of the 1rst three clusters described above and search for similar structures within the PDB database. In order to focus on the loop structure, rather than on stem variability, we consider the tetraloop and the two closing base pairs for the search (residues 4-11 in Fig.1A). The search is performed against all RNA-containing structures in the PDB database (retrieved May 4th, 2018, resolution 3.5Å or better). The database considered here consists of 3067 X-ray, 652 NMR and 177 cryo electron-microscopy (EM) structures. Note that the search is purely based on the geometrical arrangement of nucleobases, without restriction on the sequence, a particular feature that is also enabled by the use of eRMSD.

**Figure 4.**
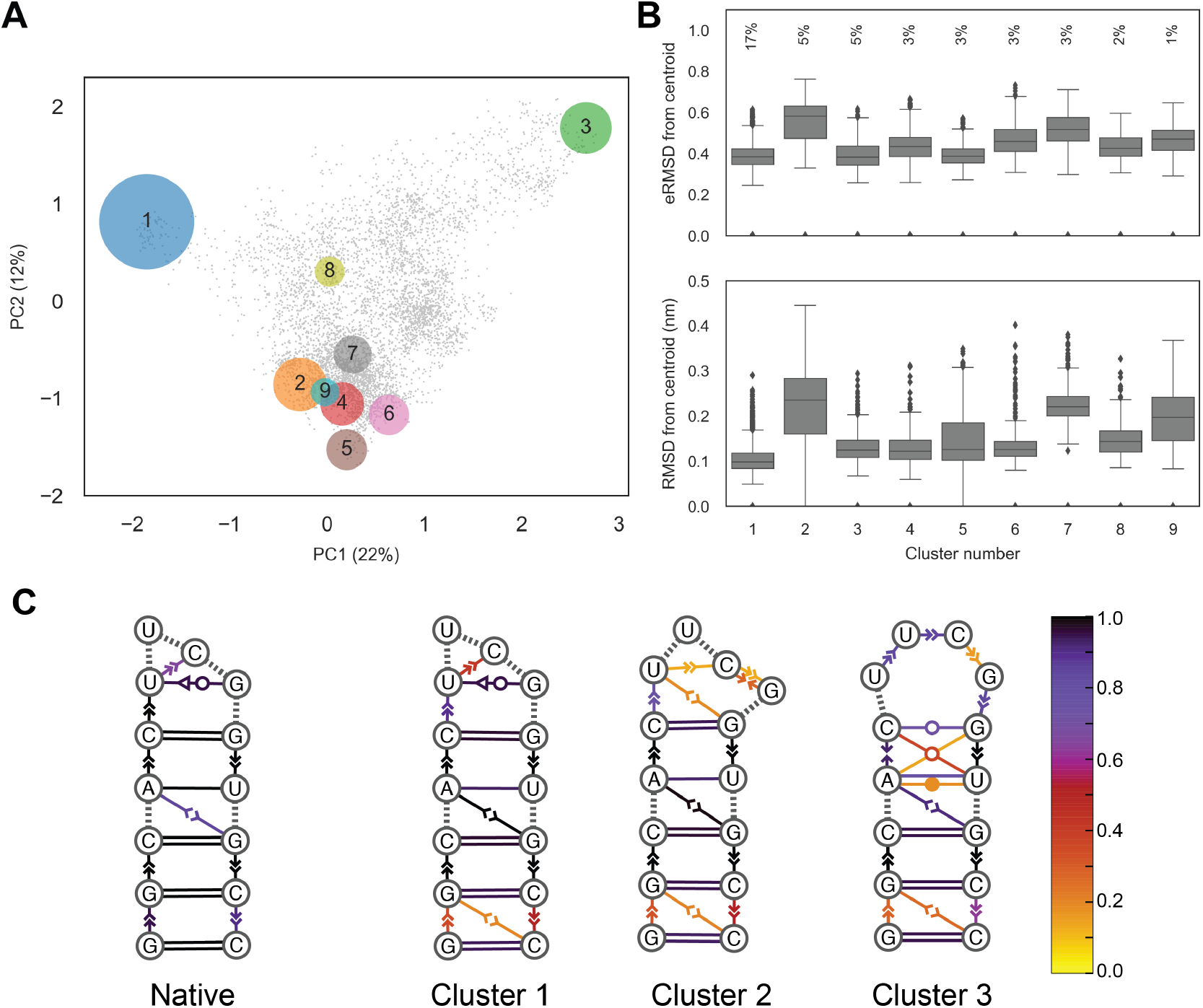
Example of a cluster analysis on the UUCG stem-loop trajectory. **A**) principal component analysis on the collection of G-vectors ***Bottaro and Lindorff-Larsen*** (***2017***). Each circle corresponds to a cluster, gray dots show unassigned structures. Circles are centered in the centroid positions, and the radii are proportional to the square root of the population. The percentage of explained variance of the first two components is indicated on the axes. **B**) Box-plots reporting eRMSD (top) and RMSD (bottom) from cluster centroids. Lower/upper hinges correspond to the first and third quartiles, while whiskers indicate lowest/highest data within 1.5 interquartile range. Data beyond the end of the whiskers are shown individually. The percentages indicate the cluster population. **C)** Dynamic secondary structure representation of the 20 native NMR conformers (PDB 2KOC) and of the first three clusters. The extended secondary structure annotation follows the Leontis-Westhof classification. The color scheme shows the fraction of frames within a cluster for which the interaction is formed.

**Figure 5.**
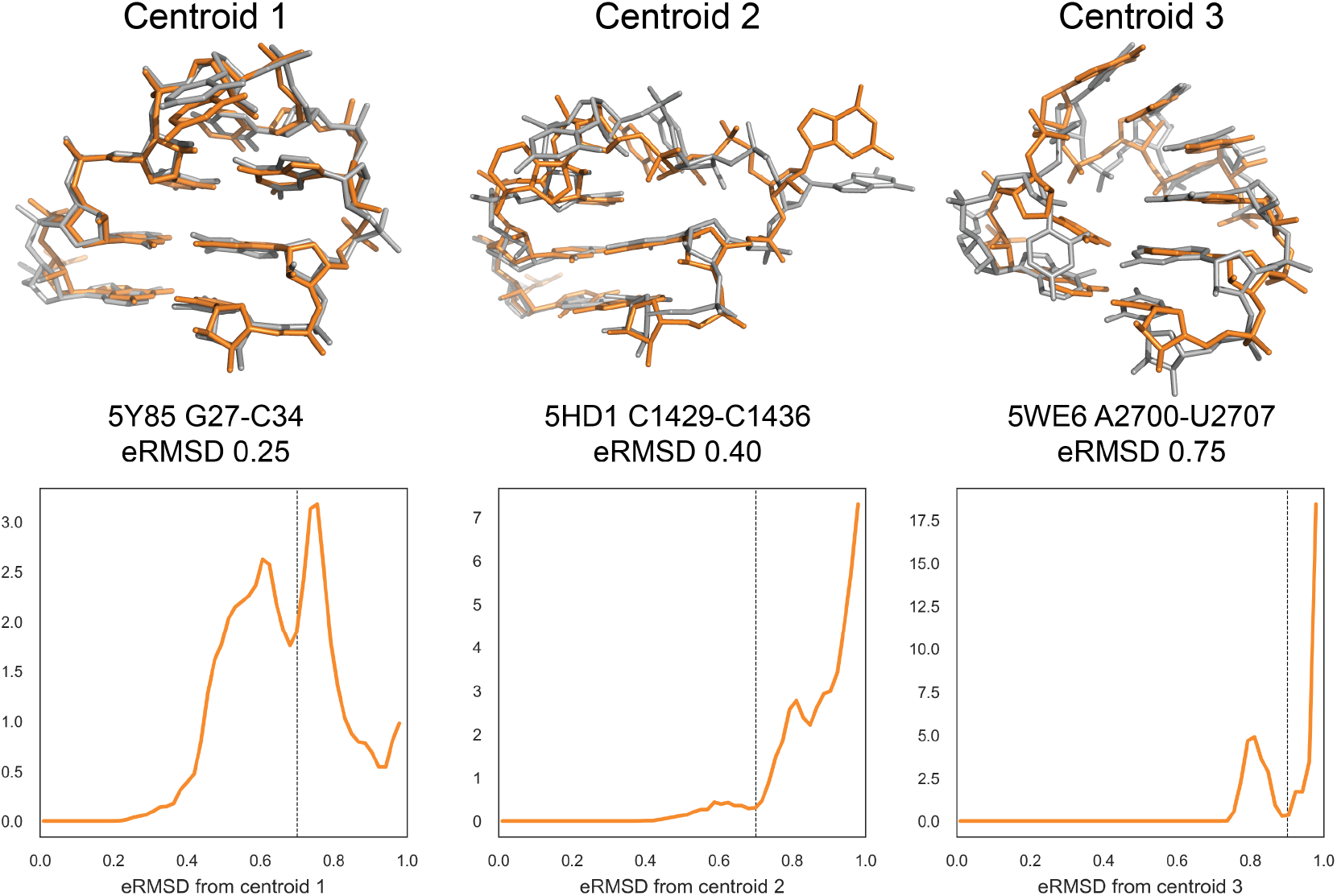
Motif search in PDB database. Top panels: centroids of the first three clusters (in gray) superimposed on the closest structures from the PDB database (orange). eRMSD between centroid and the best match are indicated, together with the associated PDB code. Bottom panels: eRMSD distribution between centroid and substructures from PDB database. Note that different distributions are obtained for different clusters, meaning that the eRMSD threshold varies depending on the motif. Distances larger than eRMSD=1 are not reported. The eRMSD threshold at 0.7 (centroids 1, 2) and 0.9 (centroid 3) is indicated as a dashed line.

Figure 5 shows the cluster centroids (gray) and the closest motif match, i.e. the lowest eRMSD substructure in the PDB database (orange). The eRMSD between the cluster centroid and the best match are indicated, together with the associated PDB code. Centroid 1 corresponds to the canonical UUCG tetraloop structure, with the signature tSW interaction between U6-G9 and G9 in syn conformation. Note that the eRMSD between centroid and best match is small (0.25), indicating that simulated and experimental structures are highly similar. Cluster 2 corresponds to a structure in which the stem is formed, C8 is stacked on top of U6 and G9 is bulged out. Centroid 3 features four consecutive stacking between U6-U7-C8-G9. Note that this latter structure is remarkably similar to the 4-stack loop described in ***Bottaro and Lindorff-Larsen*** (***2017***).

As a rule of thumb, we consider as significant matches structures below 0.7 eRMSD, but there are cases in which it is worth considering structures in the 0.7-1.0 eRMSD range as well. More generally, it is useful to consider the histogram of all fragments with eRMSD below 1, as shown in Fig. 5, bottom panels. This type of analysis makes it possible to identify a good threshold value, in correspondence to minima in the probability distributions. For example, there are no structures in the PDB with eRMSD lower than 0.7 for centroid 3. In this case, a value of 0.9 should be used instead.

In this example we performed a simple search of a structure from simulation against experimentally-derived structures in the PDB database. In Barnaba, any arbitrary motif can be used as a query by providing a coordinate file with at least the position of C2,C4 and C6 atoms for each nucleotide. Searches with more complex motifs composed by two strands (e.g. K-turns, sarcin-ricin motifs, etc.) are also possible (SM8). Additionally, Barnaba allows for inserted bases, thereby identifying structural motifs with one or more bulged-out bases.

### Elastic Network Models

Elastic Network Models (ENMs) are minimal computational models able to capture the dynamics of macromolecules at a small computational cost. They assume that the system can be represented as a set of beads connected by harmonic springs, each having rest length equal to the distance between the two beads it connects, in a reference structure (usually, an experimental structure from the PDB). First introduced to analyze protein dynamics ***Tirion*** (***1996***), ENMs are also applicable to structured RNA molecules ***Bahar and Jernigan*** (***1998***); ***Setny and Zacharias*** (***2013***); ***Zimmermann and Jernigan*** (***2014***). Barnaba contains routines to construct ENM of nucleic acids and proteins, and, as unique feature, makes it possible to calculate fluctuations between consecutive C2-C2 atoms. In a previous work ***Pinamonti et al.*** (***2015***), we have shown this quantity to correlate with 2exibility measurements performed with selective 2-hydroxyl acylation analyzed by primer extension (SHAPE) experiments ***Merino et al.*** (***2005***). Here, we show an example of ENM analysis on two RNA molecules: the 174-nucleotide sensing domain of the Thermotoga maritima lysine riboswitch (PDB ID: 3DIG), and the Escherichia coli 5S rRNA (PDB ID: 1C2X). We construct an all-atom ENM (AA-ENM), where each heavy atom is a bead, together with a cutoff radius of 7 Å. In Fig. 6 we show the flexibility of the RNA molecules as predicted by the ENM (black), that can be qualitatively compared with the measured SHAPE reactivity ***Hajdin et al.*** (***2013***) (orange).

**Figure 6.**
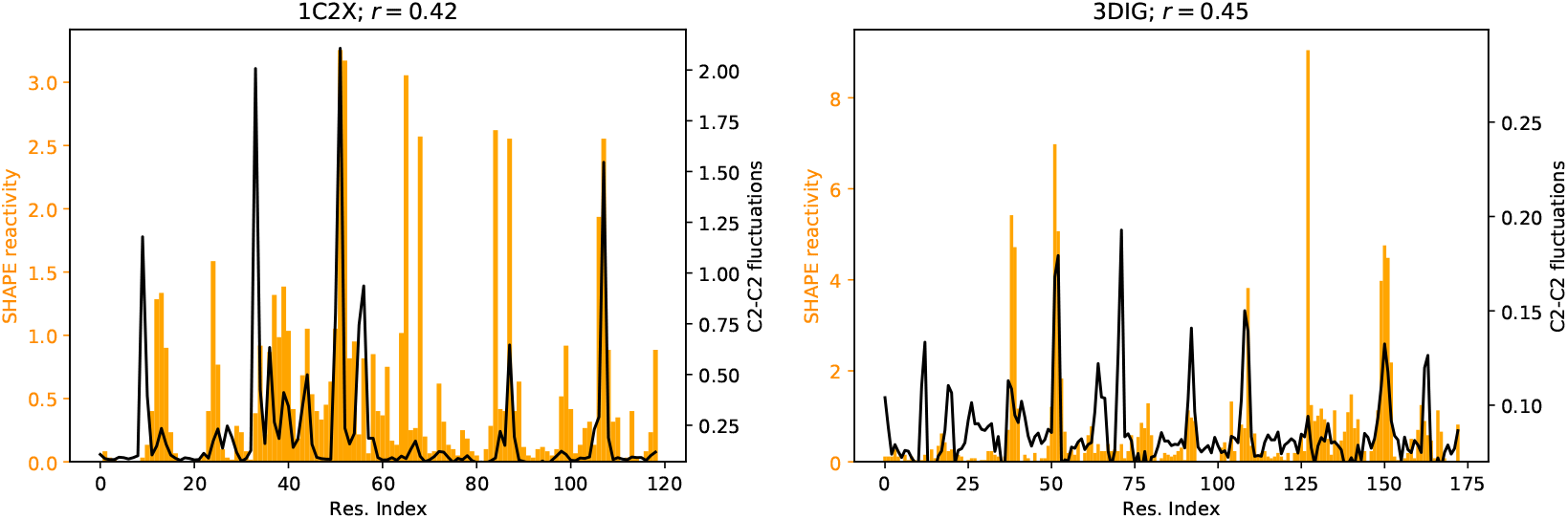
C2-C2 fluctuations as predicted by the ENM of Lysine riboswitch (right panel) and 5S rRNA (left panel). SHAPE reactivity data from ***Hajdin et al.*** (***2013***) are shown for comparison. Pearson correlation coefficient *r* between SHAPE data and ENM-predicted fluctuations is also indicated.

The implementation of the ENM in Barnaba employs the sparse matrix package available in Scipy, that allows for significant speed-ups compared to the dense-matrix implementation. Fig. 7 shows the execution time for constructing ENMs of biomolecules with sizes ranging from a few tens to several hundreds nucleotides. Calculations were performed running Barnaba on a personal computer. This, combined with the significant memory saving granted by sparse matrices representation, makes it possible to easily compute the vibrational modes and the local flexibility of large RNA systems such as ribosomal structures using a limited amount of computer resources.

## Discussion

Many RNA molecules are highly dynamical entities that undergo conformational rearrangements during function. For this reason, it is becoming increasingly important to develop tools to analyze not only single structures, but also trajectories (ensembles) obtained from molecular simulations. In this paper we introduce a software to facilitate the analysis of nucleic acids simulations. The program, called Barnaba, is available both as a Python library as well as a command line tool. The output of the program is such that it can be easily used to calculate averages and probability distributions, or conveniently used as input to the many existing plotting and analysis libraries (e.g. Matplotlib, SKlearn) available in Python.

Barnaba consists of a number of functions: some of them implement standard calculations (RMSD, torsion angles, base-pairs and base-stacking detection). A unique feature of Barnaba is the possibility to calculate the eRMSD. This metric has been successfully employed in several contexts: for analyzing MD simulations ***Kuhrova et al.*** (***2016***), as a biased collective variable in enhanced sampling simula tions ***Bottaro et al.*** (***2016a***); ***Yang et al.*** (***2017***); ***Poblete et al.*** (***2018***), to construct Markov State models ***Pinamonti et al.*** (***2017***) and to cluster RNA tetraloop structures ***Bottaro and Lindorff-Larsen*** (***2017***). In this paper we show the usefulness of this metric to monitor simulations over time, to perform cluster analysis and to search for structural motifs within trajectories/structures. This last feature can be extremely useful to experimental structural biologists, as it makes it possible to eZciently search for arbitrary query motifs within the entire PDB database. For analyzing simulations and clusters, we have found it useful to introduce a dynamic secondary structure representation, that recapitulates the variability of base-pair and base-stacking interactions within an ensemble.

**Figure 7.**
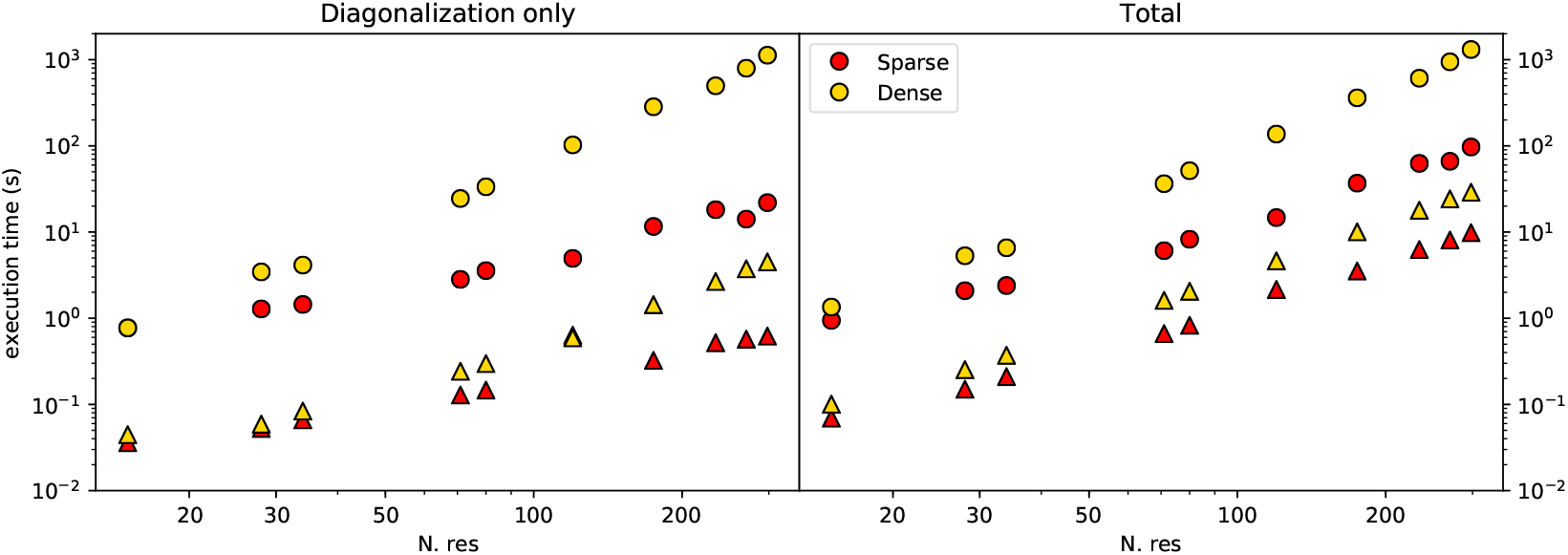
Execution time for the ENM calculation using sparse matrices (yellow) or dense matrices (red) on a 2.3 GHz Dual-Core Intel Core i5 processor, as a function of the number of residues in the RNA molecule. Results are shown both for sugar-base-phosphate (SBP) ENM (triangles) and all-atom-ENM (AA-ENM) (circles), as defined in ***Pinamonti et al.*** (***2015***). Left panel shows the time for the interaction matrix diagonalization only, right panel shows the total time including the calculation of C2-C2 fluctuations.

Another important feature of Barnaba is the possibility to back-calculate ^3^*J* scalar couplings from structures. This calculation is *per se* extremely simple. However, it can be difficult to obtain from the literature the different sets of Karplus parameters, and the calculation of the corresponding dihedral angles is error-prone.

Finally, Barnaba contains a routine to construct ENMs of nucleic acid and protein systems and complexes. This is a useful, fast and computationally cheap tool to predict the local dynamical properties of biomolecules, as well as the chain 2exibility of RNA molecules.

## Methods and Materials

### Implementation and availability

Barnaba is a Python library and command line tool. It requires Python 2.7 or > 3.3, Numpy, and Scipy libraries. Additionally, Barnaba requires MDTraj (http://mdtraj.org/) for manipulating structures and trajectories. Source code is freely available at https://github.com/srnas/barnaba under GNU GPLv3 license. The github repository contains documentation as well as a set of examples.

**Figure 8.**
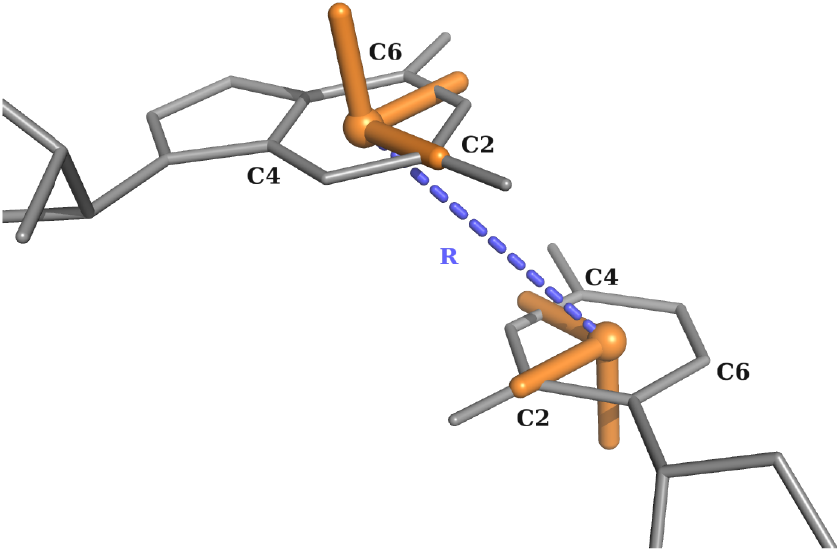
Definition of the local coordinate systems and of the vector **R** for purines and pyrimidines.

### Relative position and orientation of nucleobases

For each nucleotide, a local coordinate system is set up in the center of C2, C4, and C6 atoms. The x-axis points toward the C2 atom, and the y-axis in the direction of C4 (C/U) or C6 (A/G). The origin of the coordinates of nucleobase *j* in the reference system constructed on base *i* is the vector **R_ij_** = {*x_ij_, y_ij_, z_ij_*}. Note that |**R_ij_| = |R_ji_|** but **≠ R_ij_** is central in the definition of the eRMSD metric and of the annotation strategy described below.

### eRMSD

The eRMSD is a contact-map based distance, with the addition of a number of features that make it suitable for the comparison of nucleic acids structures. We brie2y describe here the procedure, originally introduced in ***Bottaro et al.*** (***2014***). Given a three-dimensional structure *α*, one calculates 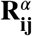 for all pairs of bases in a molecule. The position vectors are then rescaled as follows:

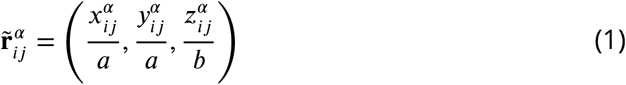

with *a* = 5Å and *b* = 3Å. The rescaling effectively introduces an ellipsoidal anisotropy that is peculiar to base-base interactions. Given two structures, *α* and *β* consisting of *N* residues, the eRMSD is calculated as

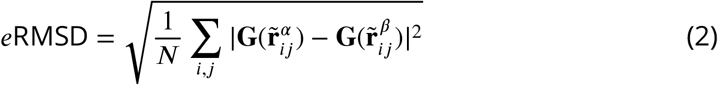

**G** is a non-linear function of **r̃** defined as:

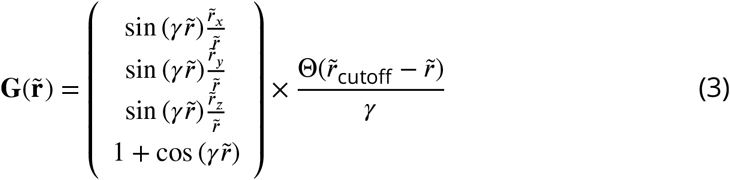

where *γ* = *π*/*r&#0303;* _cutoff_ and Θ is the Heaviside step function. Note that the function **G** has the following desirable properties:

1. |**G(r̃^α^) – G(r̃^β^)| ≈ |r̃^α^ – r̃^β^|** if ***r̃^α^*, *r̃^β^* ⪡ *r̃*** _cutoff_.
2. |**G(r̃^α^) – G(r̃^β^)| =** 0 if ***r̃^α^***, ***r̃^β^*** ≥ ***r̃***_cutoff_.
3. **G(r̃)** is a continuous function.

The cutoff value is set to *r̃* _cutoff_ = 2.4.

### Annotation

A pair of bases *i* and *j* is considered for annotation only if |**r̃**_*ij*_| *<* 1.7 and |**r̃**_*ji*_| *<* 1.7.

#### Stacking

The criteria for base-stacking are the following:

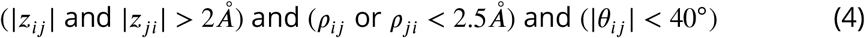

Here, 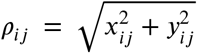 and *θ_ij_* is the angle between the vectors normal to the planes of the two bases. Similarly to other annotation approaches ***Gendron et al.*** (***2001***); ***Sarver et al.*** (***2008***); ***Waleń et al.*** (***2014***), we identify four different classes of stacking interactions according to the sign of the z coordinates:

- upward: (>> or 3’-5’) if *z_ij_ >* 0 and *z_ji_ <* 0
- downward: (<< or 5’-3’) if *z_ij_ <* 0 and *z_ji_ >* 0
- outward: (*<>* or 5’-5’) if *z_ij_ <* 0 and *z_ji_ <* 0
- inward: (*><* or 3’-3’) if *z_ij_ >* 0 and *z_ji_ >* 0

We notice that, with this choice, consecutive base pairs with alternating purines and pyrimidines result in a cross-strand outward stacking (see, e.g., Fig. 1A).

#### Base-pairing

Base-pairs are classified according to the Leontis-Westhof nomenclature ***Leontis and Westhof*** (***2001***), based on the observation that hydrogen bounding RNA bases involve three distinct edges: Watson-Crick (W), Hoogsteeen (H), and sugar (S). An additional distinction is made according to the orientation with respect to the glycosydic bonds, in cis (c) or trans (t) orientation. In Barnaba, all non-stacked bases are considered base-paired if |*θ_ij_*| *<* 60° and there exists at least one hydrogen bond, calculated as the number of donor-acceptor pairs with distance *<* 3.3*Å*. Edges are defined according to the value of the angle *Ψ* = arctan2(*y&#0302;_ij_, x̂_ij_*).

- Watson-Crick edge (W): 0.16 < *Ψ* ≤ 2.0*rad*
- Hoogsteen edge (H): 2.0 *< Ψ*≤ 4.0*rad*.
- Sugar edge (S): *Ψ >* 4.0*rad, Ψ* ≤ 0.16*rad*

**Figure 9.**
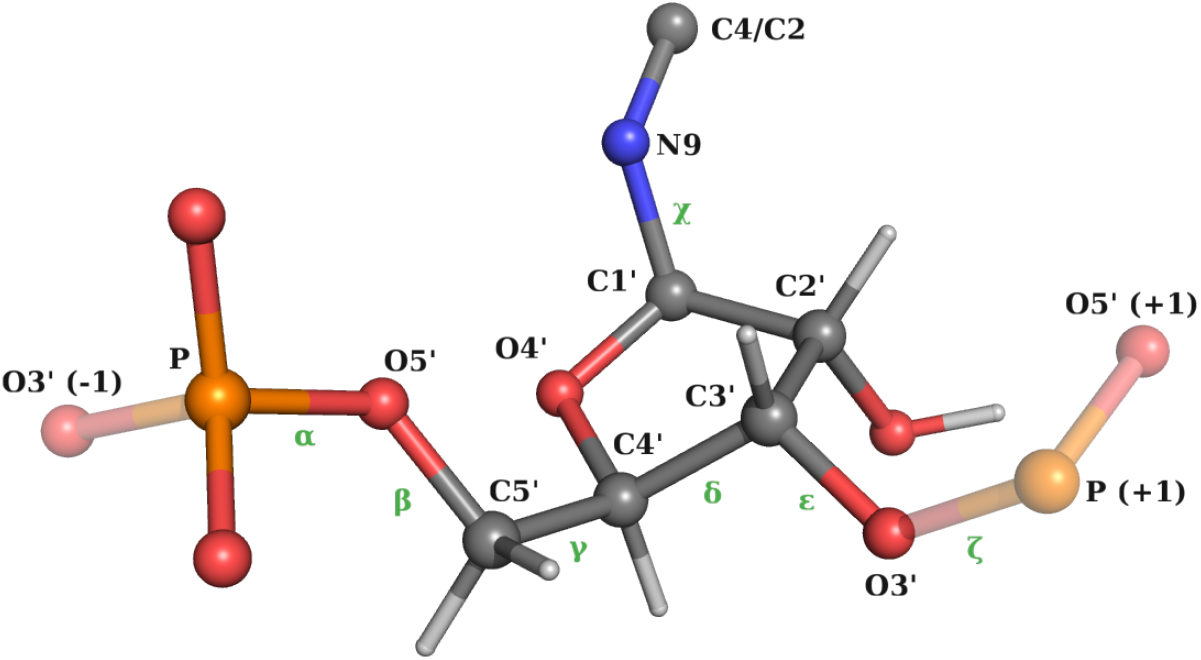
Definition of the backbone/glycosidic angles *χ****Frellsen et al.*** (***2009***).

These threshold values are obtained by considering the empirical distribution of base-base interactions shown in SM3 and discussed in Fig. 2 of Ref. ***Bottaro et al.*** (***2014***). Cis/trans orientation is calculated according to the value of the dihedral angle defined by *C*1′_*i*_ − *N*1/*N*9_*i*_− *N*1/*N*9_*j*_− *C*1′_*j′*_ where N1/N9 is used for pyrimidines and purines, respectively.

We note that the annotation provided by Barnaba might fail in detecting some interactions, and sometimes differs from other programs. This is due to the fact that for non-Watson-Crick and stacking interactions it is not trivial to define a set of criteria for a rigorous discrete classification ***Waleń et al.*** (***2014***). Typically, these criteria are calibrated to work well for high-resolution structures, but they are not always suitable to describe nearly-formed interactions often observed in molecular simulations.

### Torsion angles and ^3^*J* scalar couplings

We use the standard definition of backbone angles, glycosidic *χ* angle (O4’-C1’-N9-C4 atoms for A/G, O4’-C1’-N1-C2 for C/U) and sugar torsion angles (*v*_0_ ⋯ *v*_4_) as shown in Figures 9 and 10 ***Saenger*** (***2013***).

Pseudorotation sugar parameters amplitude *tm* and phase *P* are calculated as described in ***Rao et al.*** (***1981***)

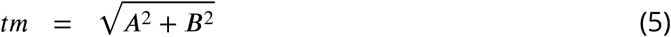

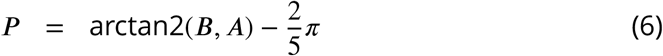

**Figure 10.**
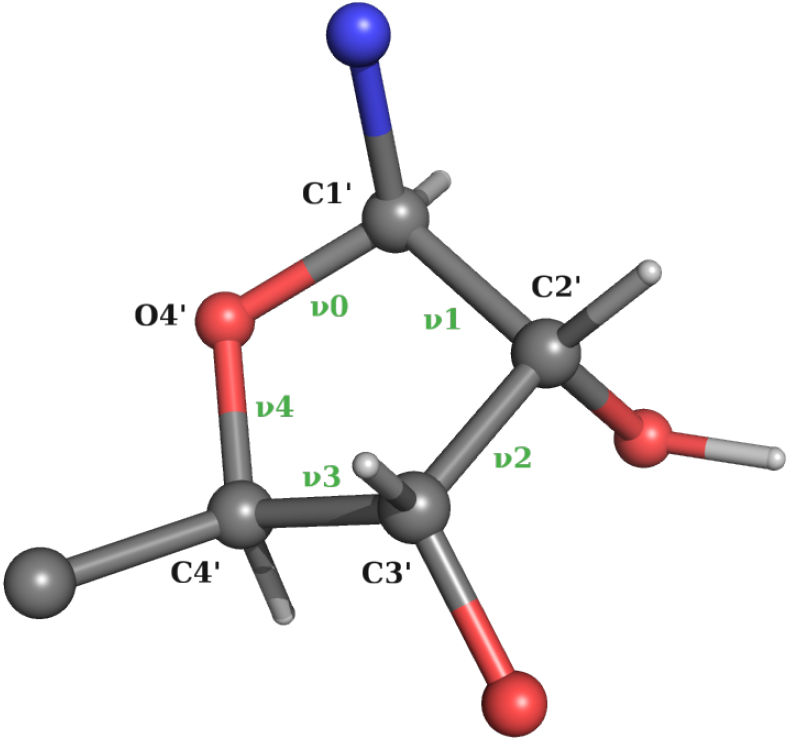
Definition of pucker angles *v*_0_ ⋯ *v*_4_

Where

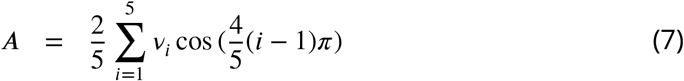

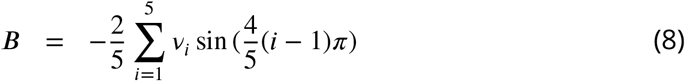

Optionally, it is also possible to calculate pseudorotation parameters using the Altona-Sundaralingam treatment ***Altona and Sundaralingam*** (***1972***)

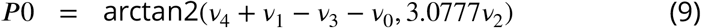

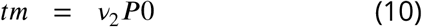

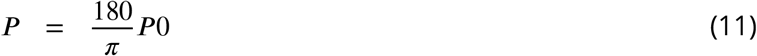

^3^*J* Scalar couplings are calculated using the Karplus equations

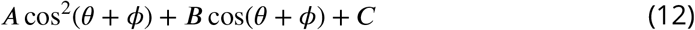

Karplus parameters relative to the different scalar couplings are reported in Table 1.

### Elastic Network Model

In ENMs, a set of *N* beads connected by pairwise harmonic springs penalize deviations of inter-bead distances from their reference values. Spring constants are set to a constant value *k* whenever the reference distance between the two beads is smaller than an interaction cutoff (*R_c_*), and set to zero otherwise. Under these assumptions, the potential energy of the system can be approximated as

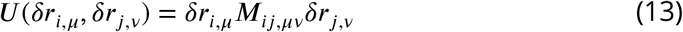

**Table 1.** Karplus parameters used in Barnaba

| Name | $\theta$ | A [Hz] | B [Hz] | C [Hz] | $\phi$ [rad] | Ref |
| --- | --- | --- | --- | --- | --- | --- |
| H1'-H2' | H1'-C1'-C2'-H2' | 9.67 | -2.03 | 0 | 0 | *Condon et al. (2015)* |
| H2'-H3' | H2'-C2'-C3'-H3' | 9.67 | -2.03 | 0 | 0 | *Condon et al. (2015)* |
| H3'-H4' | H3'-C3'-C4'-H4' | 9.67 | -2.03 | 0 | 0 | *Condon et al. (2015)* |
| H5'-P | $\beta$ | 15.3 | -6.1 | 1.6 | $-2/3\pi$ | *Lankhorst et al. (1984)* |
| H5''-P | $\beta$ | 15.3 | -6.1 | 1.6 | $2/3\pi$ | *Lankhorst et al. (1984)* |
| C4-P | $\beta$ | 6.9 | -3.4 | 0.7 | 0.0 | *Marino et al. (1999)* |
| H4'-H5' | $\gamma$ | 9.7 | -1.8 | 0.0 | $-2/3\pi$ | *Davies (1978)* |
| H4'-H5'' | $\gamma$ | 9.7 | -1.8 | 0.0 | 0.0 | *Davies (1978)* |
| H3-P(+1) | $\epsilon$ | 15.3 | -6.1 | 1.6 | $2/3\pi$ | *Lankhorst et al. (1984)* |
| C4-P(+1) | $\epsilon$ | 6.9 | -3.4 | 0.7 | 0.0 | *Marino et al. (1999)* |
| H1'-C8/C6 | $\chi$ | 4.5 | -0.6 | 0.1 | $-\pi/3$ | *Ippel et al. (1996)* |
| H1'-C4/C2 | $\chi$ | 4.7 | 2.3 | 0.1 | $-\pi/3$ | *Ippel et al. (1996)* |

where **M** is the symmetric 3*N* × 3*N* interaction matrix, and *δ***r**_*i*_is the deviation of bead *i* from its position in the reference structure.

The user can select different atoms to be used as beads in the construction of the model. The optimal value of the parameter *R_c_* depends on this choice, as described in Ref. ***Pinamonti et al.*** (***2015***).

The covariance matrix is computed as

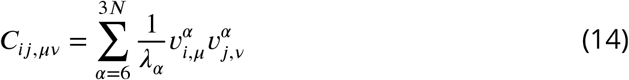

Where *λ_α_* and **v**^*α*^are the eigenvalues and the eigenvectors of the interaction matrix ***M***, respectively. The sum on *α* runs over all non-null modes of the system.

Mean square fluctuation (MSF) of residue *i* is calculated as:

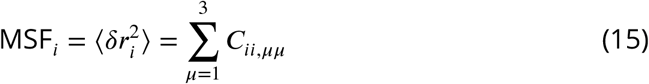

The variance of the distance between two beads can be directly obtained from the covariance matrix in the linear perturbation regime as

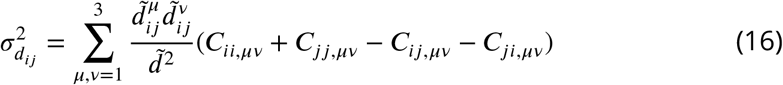

where 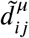 is the *μ* Cartesian component of the reference distance between bead *i* and *j*.

For most practical applications of ENMs only the high-amplitude modes, i.e. those with the smallest eigenvalues, provide interesting dynamical information. The calculation of C2-C2 distance fluctuations using Eq. 16 requires the knowledge of all eigenvectors. This can be performed by reducing the system to the “effective interaction matrix” 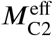 relative to the beads of interest ***Zen et al.*** (***2008***).

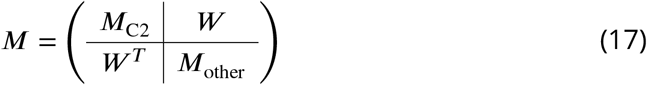

Where *M*_C2_ (*M*_other_) is formed by the rows and columns of *M* relative to the (non) C2 beads, while *W* represent the interactions between C2 and non-C2 beads. The effective interaction matrix is defined as

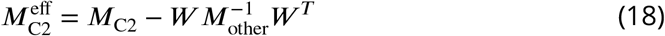

This can be computed efficiently using sparse matrix-vector multiplication algorithms. The resulting effective matrix 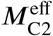 has reduced size: 1/3 for sugar-base-phosphate (SBP), 1/20 for all-atom (AA), making its pseudo-inversion considerably faster. Note that, in case one is interested in computing the C2-C2 2uctuations for a portion of the molecule only, the algorithm could be further optimized by directly computing the effective interactions matrix associated to the required C2-C2 pairs.

## Acknowledgments

We thank D.E Shaw Research for providing the simulation of the UUCG tetraloop. The research is funded by a grant from The Velux Foundations (S.B. and K.L.-L.), a Hallas-Møller Stipend from the Novo Nordisk Foundation (K.L.-L.), and the Lundbeck Foundation BRAINSTRUC initiative (K.L.-L.). G.B.,S.R, S.B and G.P. have received funding from the European Research Council (ERC) under the European Union’s Seventh Framework Programme (FP/2007-2013)/ERC grant agreement no. 306662 (S-RNA-S). W.B. is funded from VILLUM FONDEN (VKR023445) and the Danish Council for Independent Research (DFF-4181-00344).

